# PathFlow-MixMatch for Whole Slide Image Registration: An Investigation of a Segment-Based Scalable Image Registration Method

**DOI:** 10.1101/2020.03.22.002402

**Authors:** Joshua J. Levy, Christopher R. Jackson, Christian C. Haudenschild, Brock C. Christensen, Louis J. Vaickus

**Affiliations:** Program in Quantitative Biomedical Sciences, Geisel School of Medicine at Dartmouth, Lebanon, NH 03756; Emerging Diagnostic and Investigative Technologies, Clinical Genomics and Advanced Technologies, Department of Pathology and Laboratory Medicine, Dartmouth Hitchcock Medical Center, Lebanon, NH 03756; Department of Epidemiology, Geisel School of Medicine at Dartmouth, Lebanon, NH 03756

## Abstract

Image registration involves finding the best alignment between different images of the same object. In these tasks, the object in question is viewed differently in each of the images (e.g. different rotation or light conditions, etc.). In digital pathology, image registration aligns correspondent regions of tissue from different stereotactic viewpoints (e.g. subsequent deeper sections of the same tissue). These comparisons are important for histological analysis and can facilitate previously unavailable manipulations, such as 3D tissue reconstruction and cell-level alignment of immunohistochemical (IHC) and special stains. Several benchmarks have been established for evaluating image registration techniques for histological tissue; however, little work has evaluated the impact of scaling registration techniques to Giga-Pixel Whole Slide Images (WSI), which are large enough for significant memory limitations, and contain recurrent patterns and deformations that hinder traditional alignment algorithms. Furthermore, as tissue sections often contain multiple, discrete, smaller tissue fragments, it is unnecessary to align an entire image when the bulk of the image is background whitespace and tissue fragments’ orientations are often agnostic of each other. We present a methodology for circumventing large-scale image registration issues in histopathology and accompanying software. By removing background pixels, parsing the slide into discrete tissue segments, and matching, orienting and registering smaller segment pairs, we recovered registrations with lower Target Registration Error (TRE) when compared to utilizing the unmanipulated WSI. We tested our technique by having a pathologist annotate landmarks from 13 pairs of differently stained liver biopsy slides, performing WSI and segment-based registration techniques, and comparing overall TRE. Preliminary results demonstrate superior performance of registering segment pairs versus registering WSI (difference of median TRE of 44 pixels, p<0.001). Segment matching within WSI is an effective solution for histology image registration but requires further testing and validation to ensure its viability for stain translation and 3D histology analysis.

## Introduction

The field of digital pathology has made tremendous progress in the past decade with the advent of robust whole slide image (WSI) scanning devices and powerful open source software suites for image analysis and machine learning. Earlier techniques focused on morphometry and color thresholding with human-designed filters as means of separating histologic features of interest for characterization. Recent works have largely moved away from these rigid, predefined assumptions in favor of deep learning (DL) and artificial neural networks (ANN) based approaches ^1^. Applied to WSI, these models learn to extract important local morphological information and then aggregate many low level features in order to “learn” key higher-level features of the image in question. As these algorithms are validated and incorporated into clinical workflows^2^, standardizing these techniques and identifying robust, efficient pre-processing algorithms is vitally important.

Preliminary deep learning histopathology analyses focused largely on differentiating tissue of interest after pathologist annotation (e.g. Basal Cell Carcinoma versus Squamous Cell Carcinoma; neoplastic versus non-neoplastic tissue) ^3^. However, annotation of tissue is subjective, shows broad interobserver variability, and is often reliant on consensus-derived diagnostic criteria which are updated frequently. To circumvent issues associated with subjectively biased studies, a growing collection of works have sought to associate the slide information with the quantitative drivers of the disease pathology ^4^ (e.g. molecular alterations, clinical chemistry, survival, etc.), via corresponding immunohistochemistry stains (IHC), ^5,6^ and mutational panels of known oncological driver mutations (among others) ^7–9^. Furthermore, generative techniques have been developed to computationally translate one histological stain (e.g. Hematoxylin and Eosin, H&E) to a target stain of interest (e.g. Masson’s Trichrome). Such techniques have the potential to save time and money by obviating the need for costly chemical staining on an adjacent section^10^. There are a few studies that suggest the importance and relevance of studying the 3D architecture of the tissue for enhancing our understanding of disease processes. As more of these “voxel” based deep learning algorithms are published, 3D reconstructions from sequential 5 μm tissue sections are increasingly becoming relevant in investigations of disease biology, prognosis, and potential for cancer metastasis ^11,12^. Finally, having aligned, differentially stained slides for the pathologist to rapidly switch back and forth between (a very common task that pathologists do manually with glass slides) could improve productivity and clinical workflow for pathologists.

In order to associate Giga-Pixel hematoxylin and eosin (H&E)-stained WSI with information from stereotactic, molecular, or other staining characteristics, the digitized representations of subsequent 5 μm tissue sections must be aligned to a high degree of accuracy. Image registration techniques align two or more images of interest through transformation of the coordinates of one or more source images to the coordinate system of a target image^13^. These techniques have been successfully applied across many biomedical image domains including: X-Ray, CT scans and MRI modalities amongst many others^14^. Most image registration techniques can be recast as optimization problems where the similarity between two images is optimized after a series of affine transformations and deformations. However, algorithms differ in their calculation of similarity between images and how the image transformations are applied. Similarity can be defined as intensity-based or feature-based, where the former calculates a correlation-based pixel-wise or window-wise intensity measure between the source and target images, while the latter technique identifies objects of interest using algorithms such as SIFT (Scale Invariant Feature Transform) ^15^ or SURF (Speeded Up Robust Features) ^16^ that are correspondent between the two images and measures their total displacement. Based on how different the two images are, the algorithm introduces linear rigid transformations through the use of scaling, rotation, and translation operations, and non-rigid, or nonlinear operations that introduce more specialized transformations for matching. Various optimization approaches and heuristics have been proposed to discover the optimal alignment^17,18^, and recently, novel deep learning and GPU-accelerated approaches have been suggested^19,20^. For applications to histopathology image analysis, these techniques have been formalized into benchmarked studies and alignment challenges that begin to better address standard histopathology image registration issues^21–24^.

However, many of these methods have largely dismissed crucial aspects of WSI that may make them difficult to register, especially when seeking to implement registration systems at full clinical scale. WSI are typically giga-pixel sized images, which means they can occupy up to 50 GB of memory depending on magnification, amount of tissue present, and image complexity (e.g. large segments of highly heterogeneous tissue captured at high magnification require more storage space than small sections of homogeneous tissue captured at low magnification). Thus, most common clinical workstation computers lack the RAM to even open these images. This is complicated by the fact that most workstations also have the minimum required GPU to drive a small monitor and are unable to take advantage of modern CUDA GPU accelerated algorithms. While many existing registration architectures can process modestly-sized images (20Kpx by 20 Kpx slides, a typical size for WSI captured at 10X), few have been designed to handle maximum magnification WSI (40X) which can contain up to 16-fold more pixels. Most algorithms instead utilize down sampling techniques which degrades slide resolution and registration quality.

A large proportion of WSI are typically filled with white space/background. Even in tissue-heavy slides, background often comprises more than 50% of the total slide area. Moreover, this background space is not homogeneously white, but instead contains a random mosaic of thousands of near-white colors. Failing to account for heterogeneous background impedes the registration algorithm and can cause significant unwanted transformations and tissue artifacts as the algorithm struggles to incorporate both tissue and background alignment. This also unnecessarily adds to the computational burden. Current algorithms also assume that the image components (tissue fragments) contained within the WSI are in the same order and orientation as the matched tissue section, with negligible deformations, and that the paired tissue sections are morphometrically very similar to the target. When the same tissue section is scanned, destained, restained, and rescanned, it is generally assumed that there are no significant alterations to the tissue morphometry (e.g. through the introduction of new artifacts, distortions, tears, etc). In histopathology, the typical processing of tissue and the micrometer level precision of important features, render these assumptions invalid. Subsequent tissue sections are typically separated by 5um but this is not always the case and the distance between sections may be many multiples of 5 μm. Special stains and immunohistochemistry are not necessarily performed on sections that are reasonably close to the target H&E, especially in samples obtained from clinical practice where numerous stains may be obtained on serial tissue sections in discontinuous batches (in between which, the tissue block must often be shaved until coplanarity between the blade and block is restored a.k.a refaced). During sectioning, segments of tissue may enlarge, shrink, appear and disappear. Additionally, the process of destaining and restaining a slide, along with the removal of the cover slip, introduces deformations to the original tissue.

Our technique attempts to minimize as many of the above challenges as possible by first identifying background pixels with thresholding and breaking each WSI into segments defined by connected component analysis (e.g. identifying all independent tissue segments separated by background). Once all the independent tissue segments from the two WSI’s have been labelled and classified, compatible segments are identified by comparison of area and overall geometry and these segments are registered until the maximum homology matches are found. To test our hypothesis that matching and registering tissue segments is more accurate and pragmatic than co-registering multiple WSI, we introduce and test informatics software entitled *PF-MixMatch* on a set of paired H&E and Trichrome stained images of liver tissue and compare our results to one of the best-performing gigapixel image registration algorithm in the literature.

## Materials and Methods

### Materials

Twenty-six liver core needle biopsies were acquired at Dartmouth Hitchcock Medical Center via routine clinical operations, fixed, embedded in paraffin (FFPE), sectioned via microtome at 5 μm and adhered to slides. These slides were then stained with H&E or Trichrome via an automated process, according to departmental guidelines. Slides were then scanned, digitized and stored at 20x magnification using the Leica Aperio-AT2 scanner, and prepared for analysis by conversion to numpy (NPY) arrays.

### Methods

Here, we provide a description of our analytic workflow for *PF-MixMatch* (**Figure 1**). After acquiring matching WSI with H&E and Trichrome stains, a rotation detection module rotates the WSI if it suspects that it has been rotated at least 90 degrees with respect to the target WSI. Then background pixels are identified by color thresholding and set to pure white (RGB = [255,255,255]). Next tissue segments are identified and labeled by running a connected component analysis on the image, which identifies non-background objects covering an area of at least 1 × 10^15^ square pixels. Statistical parameters such as the area, perimeter, inertial or moment tensors, eigenvalues of the object, orientation with respect to the vertical, eccentricity, and major and minor axis lengths are calculated for each segment in both WSI. The statistics for each segment were represented as a vector of features which were transformed into a compressed representation. Compatible segments between the subject and target WSI are matched based on nearest neighbor searches of these compressed vectors. Then, using connected component masks, the segments are extracted and isolated using bounding boxes and rotated to a vertically aligned position (relative to the major axis) to save memory space and to hasten alignment. Finally, the proposed image pairs are registered to each other using a GPU-accelerated similarity-based (mean squared error, MSE, loss) registration technique, implemented using the backpropagation-based library Airlab^19^. The resultant distortion matrix for each segment pair is computed, allowing for the acquisition of the final registered image.

**Figure 1:**
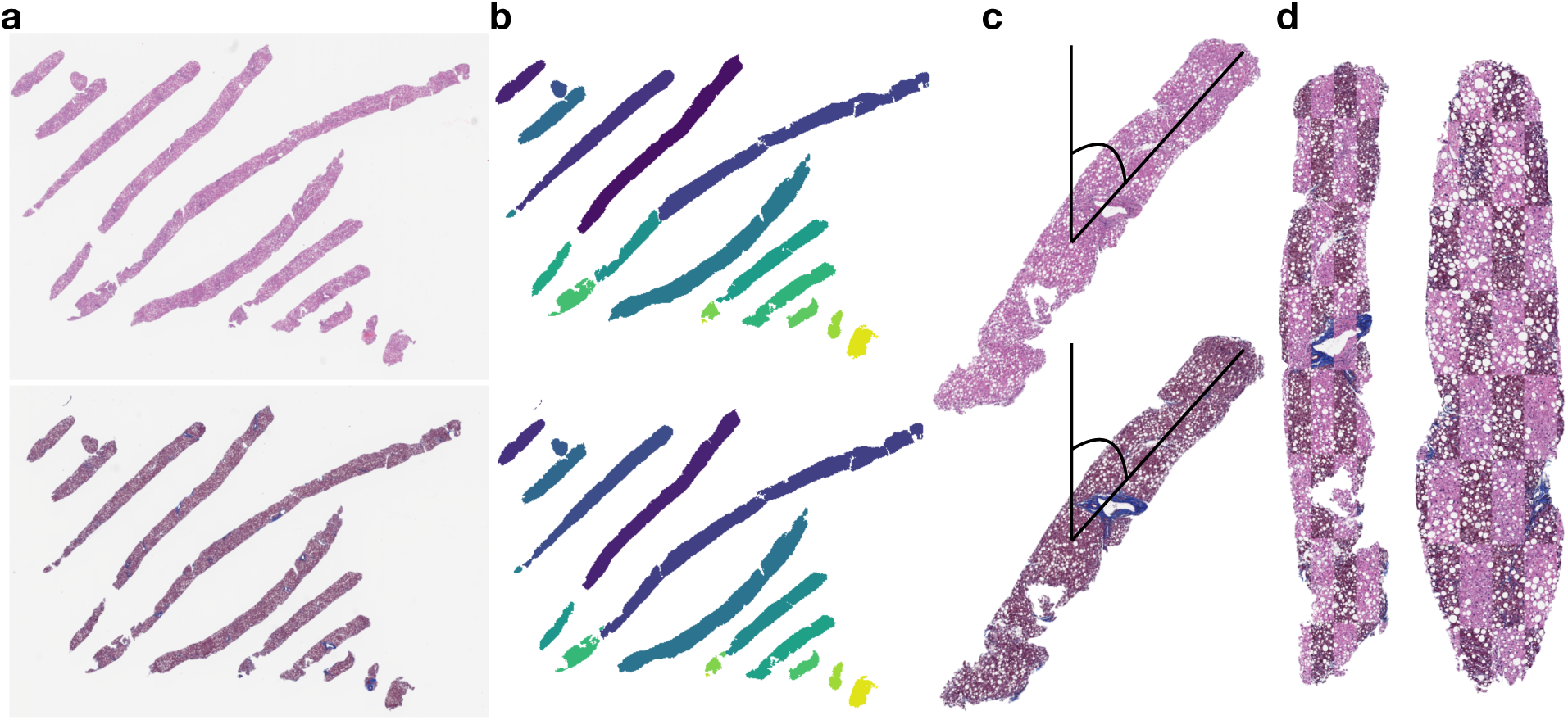
Overview of PF-MixMatch workflow: a) Paired set of images are imported and rotated to the overall correct orientation; b) Connected component analysis and shape matching identifies individual segments for further alignment; c) Individual WSI segments are rotated to the vertical position, where they await fine-tuned alignment using a registration algorithm of choice; d) Final resulting registrations of adjacent tissue sections of two segments, checkerboard pattern highlights registration alignment

### Assessment of Performance

To assess the performance of the registration method, we calculated the displacement between the corresponding marked regions on the H&E and Trichrome stains after registration, otherwise known as the Target Registration Error (TRE). As a comparison, the performance of registration on an unsegmented WSI was evaluated using a non-rigid image registration method (Reg-WSI)^17^ and compared to local segment alignment using PF-MixMatch^19^. We estimated 95% confidence intervals for the mean and median TREs across all of the morphologically distinct region for MixMatch and Reg-WSI methods using a 1000-sample nonparametric bootstrap. Due to a right-skew distribution of calculated point-wise displacements for the registrations, we conducted a Mann-Whitney U-test ^25^ to calculate whether the displacements from the Reg-WSI analysis were larger than MixMatch.

### Software Availability

We have packaged the implementation of this workflow and released it to the open-source community. The software is compatible with Python-3.6 and above, and is available on GitHub at the URL (https://github.com/jlevy44/PathFlow-MixMatch) and PyPI (*pathflow_mixmatch*) for all to download. We emphasize that this software was developed as proof-of-concept and not as production level software. We welcome additional development to address community needs through GitHub issues and offer a wiki for software usage (https://github.com/jlevy44/PathFlow-MixMatch/wiki).

## Results

### Experimental Design

Our workflow was used to register thirteen pairs of H&E stained and Trichrome stained WSI of liver tissue (randomly selected from our WSI corpus) to derive paired and registered tissue segments. Because we were not able to fit many of the original WSI into memory when using the Reg-WSI, we halved each spatial dimension of the WSI through down sampling with cubic interpolation. After registration of all WSI and image segments through the Reg-WSI and MixMatch, image and segment pairs were imported into *ASAP* ^26^ and correspondent landmark regions were identified, dotted, and annotated by a board-certified pathologist. The pathologist located the corresponding pairs of segments in their respective WSI registrations and dotted 10 or more morphologically distinctive areas in each segment. Then, for both Reg-WSI and MixMatch, we calculated and compared the registration performance as outlined in the methods.

### Registration Accuracy

The distribution of displacements was right-skewed for both algorithms (mean displacement of 843 pixels and 63 pixels for Reg-WSI and MixMatch respectively, **Figure 2**). MixMatch aligned macro-architectural slide details well, (Fig. 3a) demonstrating the ability to capture major holes, breaks, and edges in the tissue that persist between adjacent sections. The median displacement between the H&E and trichrome stains in our testing set was significantly lower for our MixMatch approach (42 pixels, 95% CI: 40px-44px, Mann-Whitney P-value <0.0001) compared with Reg-WSI (86 pixels, 95% CI: 81px-92px, and **Table 1**). In addition, out of the twelve WSI with matched segment registration data, 10 out of 12 WSIs’ Reg-WSI landmark displacements were statistically significantly greater than our MixMatch segment-based approach. For instance, Reg-WSI registration on slide 10 had physically translated entire segments of tissue to overlap with adjacent segments, causing significant registration errors (Figure 3b). Reg-WSI posted one statistically significant lower TRE than MixMatch for one of the remaining two slides. A per slide breakdown of registration errors can be found in **Table 1**.

**Table 1:** Per slide breakdown of target registration errors for the Reg-WSI and MixMatch methods; * denotes a statistically significant result with p ≤ 0.05; † denotes slide from which Reg-WSI yields statistically smaller TRE than MixMatch.

| Slide ID | Reg-WSI Mean TRE (pixels) | MixMatch Mean TRE (pixels) | Reg-WSI Median TRE (pixels) | MixMatch Median TRE (pixels) | P-Value |
| --- | --- | --- | --- | --- | --- |
| 10 | 5181 | 78 | 4931 | 60 | 5.43E-52 * |
| 144 | 97 | 76 | 83 | 58 | 0.0033 * |
| 162 | 92 | 54 | 89 | 32 | 3.52E-10 * |
| 169 | 72 | 37 | 70 | 27 | 1.64E-08 * |
| 212 | 77 | 67 | 70 | 71 | 0.229 |
| 247 | 54 | 35 | 57 | 25 | 0.00251 * |
| 249_1 | 90 | 46 | 80 | 35 | 2.44E-17 * |
| 43 | 290 | 174 | 104 | 144 | 0.976 † |
| 51 | 81 | 39 | 76 | 35 | 0.000107 * |
| 71 | 48 | 26 | 53 | 13 | 0.00288 * |
| 86 | 86 | 57 | 72 | 46 | 1.84E-09 * |
| 99 | 157 | 36 | 89 | 38 | 0.00099 * |
| Overall | 843 | 63 | 86 | 42 | 2.01E-72 * |

**Figure 2:**
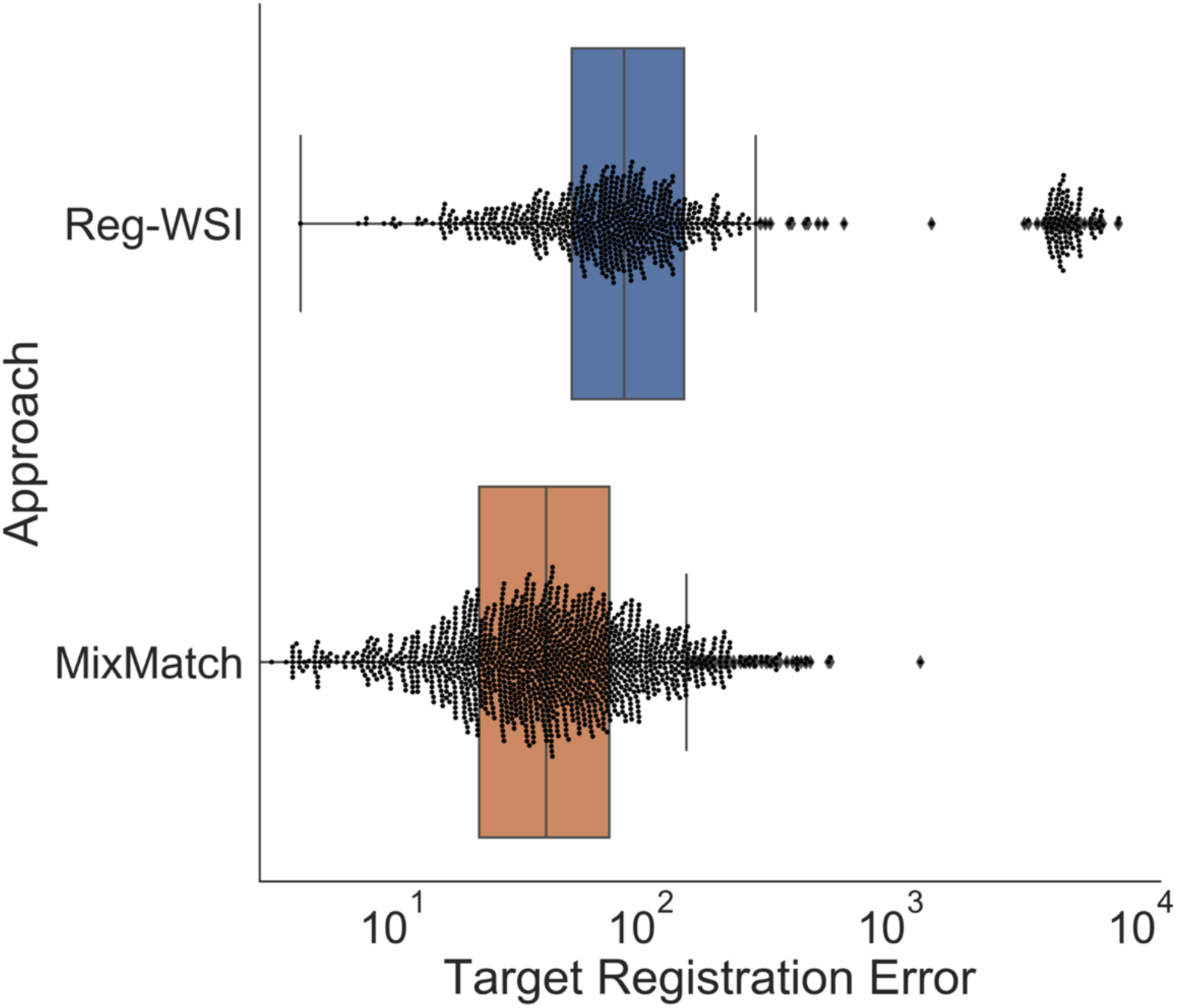
Swarm plot embedded in box and whisker plot of target registration errors for distinct morphological regions identified on the WSIs (n=766 regions) and segments (n=1365 regions)

**Figure 3:**
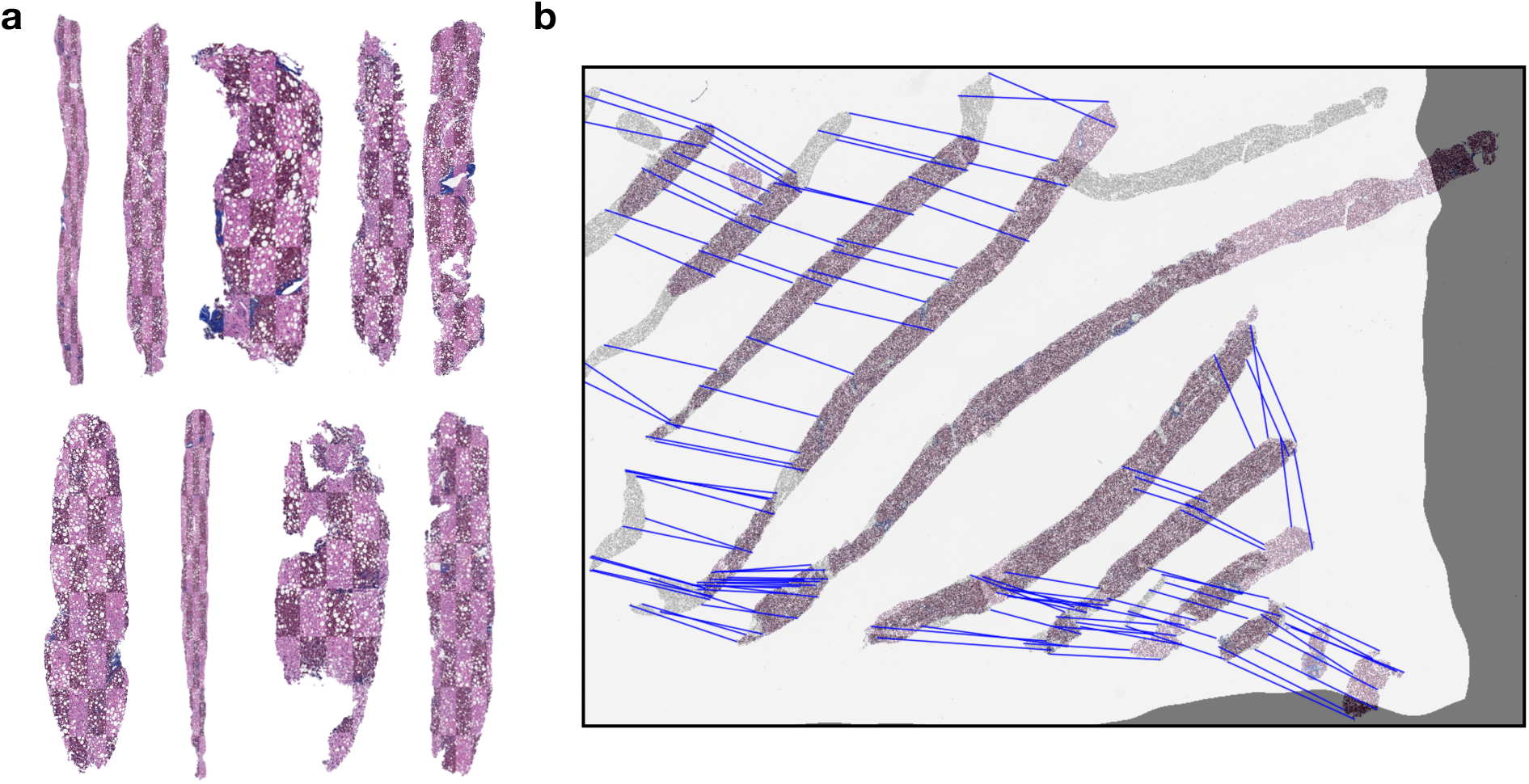
Visual comparison of registrations obtained for slide ID 10 using approaches: a) MixMatch, b) Reg-WSI

To test whether the additional Reg-WSI registration errors could be attributed to heterogeneous background alone, we performed the same background deletion step as in MixMatch on each WSI using connected component analyses and reran Reg-WSI on these background deleted WSI; as compared to the MixMatch pipeline, Reg-WSI still had larger registration error even without heterogeneous background (p=3e-10).

## Discussion

Entire gigapixel WSI images can be difficult to co-register because they are large, have non-matching features between adjacent sections, and are prone to tissue artifacts. Thus, we sought to investigate strategies that utilize connected components and shape matching to reduce the memory constraints of the algorithm and improve local alignment. Our results suggest that breaking WSI into discrete tissue segments and registering these segments (MixMatch) provides better histologic registration accuracy than attempting to register an entire WSI (Reg-WSI). Utilizing the MixMatch approach effectively eliminated the possibility of “accidental translation” (e.g. the algorithm overwarping an image) by matching the segments a priori and then performing the local alignment, which yielded more exact alignments (Figure 3b). Indeed, MixMatch performed better then Reg-WSI on all but two test cases.. As a consequence of performing registration on local tissue components rather than the entire WSI, down-sampling was not necessary to fit the entire slide into memory (as the tissue segments are significantly smaller than the unmanipulated WSI) achieving robust performance and memory efficiency gains compared to using other WSI registration techniques.

While MixMatch has demonstrated the ability to accurately capture correspondent macroarchitectural details between adjacent slides and reduce memory constraints by breaking the entire WSI into its respective components, there are a few caveats and limitations to our approach. While we demonstrated macroarchitectural correspondence, microarchitectural features (eg. nuclei, small holes, vacuoles, cells) may be absent or different across serial tissue layers, thus precluding analysis of cell level alignment. We did not fully address duplicated tissue sections placed side-by-side on WSI (a common practice in certain biopsy types), an issue which must be addressed prior to clinical use. Furthermore, there were instances in which connected components denoted two slightly connected tissue segments in one section as a single component, while separating these tissue segments in the adjacent section. The shapes of the resulting fragments are unable to be matched and thus warrant additional considerations such as allowing maximum homology matches within larger tissue segments, manual separation ^27^ or utilization of clustering techniques to naturally separate the tissue ^28–30 31–33^. Finally, we tested this technique on a small number of slides; further validation on a larger set may be warranted.

In this study, we did not seek to compare solely the differential accuracies of the underlying registration algorithms, but also to highlight the advantages of dividing WSI into discrete tissue segments prior to registration with an eye toward computational efficiency. The finetuning alignment technique employed by MixMatch utilized the Airlab registration library; however, it may be fruitful to investigate other tools such as HistoReg, ^18^ Recursive Cascaded Networks, ^34^ and other approaches ^35,36^ as means to better align individual segment pairs as it applies to their specific domain. Additionally, there are many registration loss functions that can be utilized to improve registration quality, such as cross-correlational, structural similarity, and mutual information losses, which may warrant further study. Future works will focus on improvements to shape matching ^37,38^, microarchitectural alignment on restained slides, finetuning registrations through patched based alignment ^39,40^, apply distributed computing techniques for high throughput processing ^41^ and investigate the impact of this histology registration on 3D macroarchitecture reconstruction of tissue and on stain translation techniques through Pix2Pix ^10,42 43 44^.

While we do not claim our software provides the optimal solution for registration in histopathology, our preliminary results indicate segment matching within WSI is an effective technique to provide high-resolution accurate GPU-accelerated histology image registrations that may be readily deployable for 3D reconstruction of tissue and for deep learning stain translation techniques. We encourage the greater community to consider employing these techniques into their pathology informatics workflows and trying their own rendition of these shape matching and alignment algorithms for further optimization.

## Authors’ Contributions

The conception and design of the study were contributed by JJL and LJV. Implementation, programming, data acquisition, and analyses were by JJL. All authors contributed to writing and editing of the manuscript.

## Acknowledgements

Not applicable.

